# A Reinforcement Learning-based Approach for Dynamic Privacy Protection in Genomic Data Sharing Beacons

**DOI:** 10.1101/2024.10.28.620587

**Authors:** Masoud Poorghaffar Aghdam, Sobhan Shukueian Tabrizi, Kerem Ayöz, Erman Ayday, Sinem Sav, A. Ercüment Çiçek

**Author notes:** Equal Contribution (Masoud Poorghaffar Aghdam, Sobhan Shukueian Tabrizi).

## Abstract

The rise of genomic sequencing has led to significant privacy concerns due to the sensitive and identifiable nature of genomic data. The Beacon Project, initiated by the Global Alliance for Genomics and Health (GA4GH), was designed to enable privacy-preserving sharing of genomic information via an online querying system. However, studies have revealed that the protocol is vulnerable to membership inference attacks, which can expose the presence of individuals in sensitive datasets. Various countermeasures, such as noise addition and query restrictions, have been proposed but are limited by static implementation, leaving them prone to attackers that can adapt and change strategies. In this study, we present the first reinforcement learning (RL)-based approach for dynamic privacy protection of the beacon protocol. We employ a multi-player RL setting where we train (i) a “Generic-Beacon-Defender” agent who can adjust the honesty rate of its responses, against (ii) a “Generic-Beacon-Attacker” agent who can choose the order of the queries and ask random queries to make the beacon think it is a regular user. This is the first defense mechanism capable of adapting its strategy in real time based on user queries, distinguishing between legitimate users and potential attackers, and applying tailored policies accordingly. By doing so, this method enhances both privacy and utility, effectively countering sophisticated and evolving threats. The code and the models are available at github.com/ciceklab/beacon-defense-strategies.

## 1 Introduction

The rise of genomic sequencing power has led to a dramatic increase in available genomic datasets. However, since an individual’s genomic data serves as a unique identifier, privacy concerns have emerged regarding the sharing of this information. The Beacon Project was launched by the Global Alliance for Genomics and Health (GA4GH) [1] to enable privacy-preserving sharing of genomic data using an online querying system. A researcher searching for specific alleles (nucleotides) can use the beacon’s online interface to query the connected datasets. For each dataset, the beacon provides a simple ‘yes’ or ‘no’ response. Beacons are usually associated with a sensitive phenotype or condition, e.g. Autism Speaks MSSNG Dataset. Consequently, inferring an individual’s membership in a particular beacon leads to privacy-sensitive information leakage. However, the protocol does not disclose allele frequencies and in theory, ensures that a positive response cannot be linked to any individual participant. The protocol facilitates cross-site collaborations by minimizing paperwork unless a relevant dataset is identified. The Beacon Network, a central system implementing this protocol, offers query access to over forty genomic databases and is widely used by the community [2].

Despite being designed to preserve privacy, several studies have demonstrated that the protocol leaks sensitive information about individuals in the dataset and is prone to membership inference attacks. Firstly, Shringarpure and Bustamante (2015) illustrated that an individual’s membership in a beacon can be inferred using a likelihood ratio test (LRT) by querying the beacon for several hundred SNPs of that individual (*SB Attack* ) [3]. Later, their work was extended by Raisaro et al. (2017) who introduced the *Optimal Attack* [4]. This attack incorporates information about Minor Allele Frequencies (MAFs) and decreases the number of needed SNPs to be queried dramatically to only a few. von Thenen et al. (2018) used linkage disequilibrium and high-order Markov chains to infer beacon responses and infer hidden SNPs of a victim to show the risk is even larger than already anticipated because these attacks can circumvent the suggested countermeasures [5]. In addition to membership inference attacks, Ayoz et al. (2021) uncovered a new vulnerability in the beacon protocol, demonstrating that it is possible to reconstruct a victim’s genome using beacon responses obtained at different time points [6].

The disclosed risks have started a new line of research on countermeasures to protect the privacy of individuals and limit information leakage over beacon responses. The goal of countermeasures is to prevent the reidentification of any individual in the beacon while preserving the system’s utility in accurately answering queries. The first method is *strategic flipping* of the beacon responses [7]. This method ranks SNPs by their differential discriminative power which is a method to find out which SNPs reveal more information regarding an individual’s membership. The responses for top-ranked SNPs are then flipped. This approach outperforms baseline strategies like random flipping. However, the system still provides incorrect answers for the most informative SNPs, significantly diminishing the utility. Raisaro et al. (2017) introduced a budget-based approach where each individual in the beacon is assigned a query budget [4]. If a registered user queries too many rare SNPs for an individual, the budget expires, and the beacon responds as if that person is not in the dataset. The authors show that this method outperforms baselines such as random flipping and responding *yes* only if at least 2 individuals have the allele. Yet, von Thenen et al. (2018) later demonstrated that this approach is ineffective when the attacker infers beacon answers without posing them, using correlations among SNPs. Ayöz et al. (2020) proposed to include the relatives of the beacon participants into the dataset as a source of noise [8]. Finally, Cho et al. (2020) incorporated differential privacy into flipping strategies to provide formal privacy guarantees. Yet, aforementioned works [7,4,9] mostly focus on a batch query setting, where all queries are submitted at once. This is an unrealistic assumption because the actual protocol works in an online setting where queries are sequentially posed and the attacker can easily change strategies upon observing the defense. Moreover, they consider a fixed risk threshold which determines individuals whose privacy is breached. This approach is also unrealistic, as it may shift as strategies evolve. Optimization-based defense methods address this by using adaptive thresholding schemes [10,11]. An adaptive setting allows parties to adjust strategies with each query, making static countermeasures ineffective and impractical. Following this, Zhang et al. propose the first game theoretic approach to protect beacon summary statistics [12]. They propose a Bayesian attack and simulate the system to find the parameters that lead to the approximate Nash equilibrium. They model both the attacker and the beacon as neural networks, training them in an adversarial setting. However, this approach learns an optimal set of parameters to perturb responses for all queries, without distinguishing whether a dataset participant is under attack or if the queries come from an attacker or a legitimate user. Thus, every user is treated as a potential attacker, which compromises the utility. Additionally, this game-theoretic approach requires exploring the entire search space of possible strategies, which is computationally intractable, as we also demonstrate in our results. As a result, relying on an approximation to the optimal solution becomes necessary. Consequently, no defense strategy currently exists to protect individual privacy against membership inference attacks that can dynamically adjust strategies based on the beacon’s actions in an online sequential game setting.

Our first contribution is formulating the interaction between the users and the beacon as a Stackelberg game which is more suitable to model the sequential interaction between the user and the beacon as opposed to the Bayesian game which assumes both parties simultaneously move. We formulate the first game theory-based defender to address LRT-based attacks. While it is effective, we show that due to the large strategy space, the defender can handle only a few queries.

Our second and most important contribution is using a reinforcement learning (RL)-based approach to search for the optimum policy for the first time. That is, we train an agent that is going to strategically handle user queries and adjust its honesty rate while responding to queries to protect the system participants from re-identification. RL algorithms offer a potential solution for beacon services to swiftly optimize their privacy policies based on the observations of the environment. This adaptability ensures that privacy policies can change in response to new threats, rather than remaining static and potentially vulnerable. Moreover, the goal-directed nature of RL in privacy preservation allows it to learn and focus on the most relevant features of the problem to avoid exploration of all possible strategies. This not only accelerates identifying effective privacy policies but also can handle larger query sets, which are typically more challenging to secure.

We introduce the *Generic-Beacon-Attacker*, GBA, and the *Generic-Beacon-Defender*, GBD, who are trained against each other in a multi-player setting. The GAA learns to pose queries in any order of MAFs and even pose irrelevant queries (e.g., SNPs that the victim does not have) to confuse the defender. The GBD learns to detect such patterns, identify an attack, and adjust its honesty level to confuse the GBA. We show that GBA can successfully preserve the privacy of the participants while maintaining the utility for regular users. Thus, this is the first study that can adapt the privacy policies in response to new threats, rather than being static and potentially vulnerable to updates on the attacker protocol. Moreover, the model does not require direct training on dataset participants’ genetic information. It only uses statistical information about the dataset such as MAFs for queries and LRT distributions in the dataset which means the training process can be outsourced without sharing the controlled genetic data. Our experiments show that this RL-driven approach outperforms static defenses, offering greater resilience to advanced attacks and stronger privacy across complex query sets. Our implementation is available at github.com/ciceklab/beacon-defense-strategies.

## 2 Methods

### 2.1 Problem Statement

Despite its privacy-preserving design, the Beacon protocol is vulnerable to membership inference attacks that can reveal whether an individual is part of a dataset, potentially compromising sensitive information. Existing defenses [7,4,12,13], aim to mitigate these risks but often assume static or batch-query environments. Additionally, current game-theoretic approaches [12,13] model the problem as a Bayesian game which provides a single perturbation-based solution for all users, leading to sub-optimal privacy protection. In reality, attackers can adapt their strategies in sequential-query settings, making these defenses less effective. Hence, we focus on the problem of designing defenses for evolving and more sophisticated attacks.

### 2.2 The System and Threat Model

The beacon system responds to received queries on the presence of alleles in the connected database(s) and return ’yes’ or ’no’ response for each dataset. Without losing generality, we assume the beacon is connected to a single database. The protocol operates in an authenticated online setting. That is the querier logs in before posing potentially multiple queries. The system has access to previous queries made by the same querier. This continuous access enables the protocol to accumulate information over time, improving its effectiveness in detecting membership inference attacks. In this context, the beacon assigns probabilities to “yes” responses, reflecting the honesty rate of the system. On the other hand, the honesty rate for “no” responses is always set to one, as research has demonstrated that altering “no” responses to “yes” does not enhance privacy protection for individuals within the system [10]. Note that the beacon publicly shares the honesty rate associated with every answer in our setting, which assumes a very strong adversary.

We consider a passive adversary who has access to an individual’s genome and seeks to perform a likelihood ratio test (LRT)-based membership inference attack. The adversary aims to conduct a hypothesis test to confidently determine whether the target individual’s genomic data is included in the dataset accessible via the beacon protocol. We assume the adversary can submit multiple queries without any auxiliary information and that the queries and corresponding responses are independent of each other, i.e., not in linkage disequilibrium. Additionally, we assume the adversary has access to the genomes of a control group of individuals with similar genetic backgrounds to individuals in the beacon dataset, denoted as *C*.

### 2.3 Overview of Our Solution

In this work, we propose a novel approach for defending against membership inference attacks in genomic beacon systems by leveraging both game theory and reinforcement learning (RL). Our solution begins by modeling the interaction between the attacker and the defender as a sequential Stackelberg game. With this, the beacon anticipates and responds to the strategies of the attacker, enabling more adaptive and dynamic defenses compared to previous static models. Unlike prior game-theoretic approaches that relied on simultaneous moves from both parties, our formulation takes into account the temporal nature of queries, allowing the defender to adjust its strategy based on real-time observations of the attacker. While effective against membership inference, this foundational model is limited by the computational burden of large strategy spaces, which we mitigate through strategic query handling.

Building on this foundation, we introduce a reinforcement learning (RL)-based solution that enhances the defender’s ability to adapt to evolving attack strategies. By training an RL agent, the system learns optimal policies for managing the trade-off between privacy preservation and system utility. The agent continuously adjusts its response strategies in real-time, modifying the honesty level of responses to queries based on observed patterns in user behavior. This dynamic approach enables the system to effectively block attackers while preserving high query accuracy for legitimate users. Additionally, the RL model learns from dataset summaries rather than individual genomic data, enabling scalable, privacy-preserving training.

### 2.4 Background on Game Theory and Reinforcement Learning (RL)

A Bayesian game is a game with simultaneous moves/actions where players have incomplete information about each other. An *α*-Bayesian game is an extension where instead of having precise probabilistic beliefs, players work with a set of possible probability distributions. The parameter *α* reflects the players’ level of ambiguity aversion. Stackelberg games, on the other hand, involve a leader-follower structure, where the leader moves first and the follower responds. The leader’s strategy accounts for the fact that the follower will optimize their response after observing the leader’s move. RL has been used to model Stackelberg games in various domains [14,15,16]. RL enables both the leader and follower to learn and adjust their strategies dynamically in real time. It is very effective in games with large actions or state spaces because it can approximate optimal strategies without exhaustively exploring all possibilities.

RL techniques rely on Markov Decision Processes (MDPs), including the following fundamental components: **State** *s* represents the state of the world; **Action** space *A* is the set of all valid actions available to a player in a given environment; **Reward** *R* function computes the reward for a specific state and action, and sometimes it also depends on the next state; **Policy** *π* is a rule used by an agent to decide what actions to take by mapping the states of the environment to actions. **The environment** conveys the current state to the agent and changes with respect to the agent’s actions by transitioning to a new state. It provides a reward or penalty; **Value Function** *V* estimates how good a state is, and *Q*^π^ estimates how good a state-action pair is in terms of long-term rewards. The difference is the **advantage** *Â*; **Discount Factor** *γ* ∈ [0, 1] weighs how much future rewards are valued compared to immediate rewards.

Similar to many membership inference attacks in the literature [3,4,5,10], we employ a LRT-based attack. The genome of an individual *i* can be represented as a binary vector, ***d***_***i***_ = {*d*_*i*1_, …, *d*_*ij*_, …, *d*_*im*_}where *d*_*ij*_ denotes the presence of SNP *j* (1 for “yes”, 0 for “no”) and *m* denotes the number of known SNPs. The beacon, after receiving a query set, *Q* consisting *k* queries, generates a binary response set ***x*** where *x*_*j*_ denotes the presence of the SNP in the beacon for the *j*^*th*^ query in *Q* (*Q*_*j*_).

Assuming that the MAFs for the SNPs are public, the optimal attacker constructs a list *Q* which contains the SNPs of the victim. It is sorted in ascending MAF order. Let the null hypothesis (*H*_0_) refer to the case in which the queried genome is not in the Beacon and the alternative (*H*_1_) is otherwise.

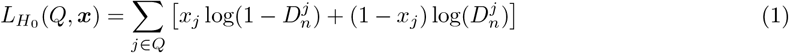

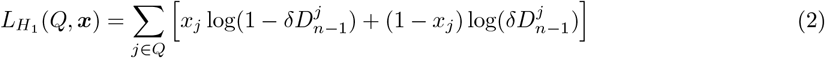

where *δ* represents the probability of sequencing error, 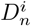 is the probability that none of the *n* individuals in the beacon having the allele at position *j* and 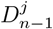 represents the probability of no individual except for the queried person having the allele at position *j*. 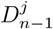 and 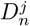 are defined as follows: 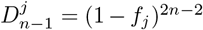 and 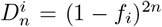 , where *f*_*j*_ represents the MAF of the SNP at position *j*. Then, the LRT statistic for person *i* in the beacon is determined as Equation 3.

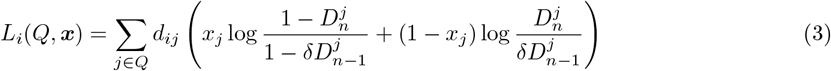

While Raisero et al. (2017) assume that the beacon is honest with all answers, Venkatesaramani et al. (2023) incorporate the fact that the beacon might be lying in the formulation. They introduce a binary variable *y*_*j*_ to indicate if the response for the *j*^*th*^query was flipped. They consider *Q*_1_ as the subset of *Q* with *x*_*j*_ = 1 and *Q*_0_ as the subset with *x*_*j*_ = 0. Let 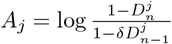 and 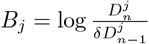 . Thus, by defining *h*_*j*_ as the complement of *y*_*j*_, representing honesty, the LRT function can be rewritten as in Equation 4.

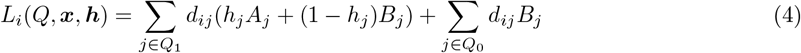

We use Equation 4 but relax the assumption that ***h*** is a binary vector. We define ***h*** as {*h*_*j*_ ∈ ℝ : 0 ≤ *h*_*j*_ ≤ 1} where *h*_*j*_ = 1 indicates an honest response and if *x*_*j*_ = 0, then *h*_*j*_ = 1. Thus, the beacon can incorporate uncertainty into the responses and fine-tune the honesty level as desired.

The adversary can decide if the individual *i* is a member of the dataset when *L*_*i*_(*Q*, ***x, h***) falls below a predefined fixed threshold *θ* [7,17]. The attacker simulates the attack on the control group *C* and picks the one that balances precision and recall. However, *fixed thresholding* can be bypassed using various defense mechanisms [10]. We rather adopt *adaptive thresholding*, which is a more effective and realistic technique that can work in an online setting and can adjust the threshold *θ* based on the response set ***x*** to the queries *Q*: *θ*(*Q*, ***x***). It ensures the false positive rate for an attack does not exceed a predefined rate [10].

### 2.5 Privacy and Utility Definitions

We define the function *p*_*i*_ to measure the privacy risk of an individual *i*. This function compares the log-likelihood ratio test (LRT) value of individual *i* with the LRT values of all individuals in a control group. It then calculates the percentage of control group members whose LRT values are greater than that of individual *i*. It is defined as 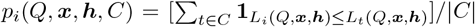 where *L* represents the LRT, computed based on the attacker’s set of queries *Q* and sharer’s responses ***x*** as previously described. This function allows us to quantify the risk of re-identification over time as more queries are posed: A lower, rapidly decreasing value is interpreted as high risk, indicating that the probability of correctly re-identifying an individual is increasing.

We quantify the utility of the beacon service using function *u*(*Q*, ***h***) = [Σ_*j*∈Q_(*h*_*j*_)]/|*Q*|. The function returns the average honesty level of the responses provided by the system which is captured by the variable ***h*** in the *LRT* function defined in Equation 4 where *h*_*j*_ represents the honesty level for the *j*^*th*^ query in query set *Q*.

### 2.6 Our Novel Defense Strategies

Here, we present a novel game theory-based defense mechanism against LRT-based attacks which can handle a few queries due to the large strategy space. Then, we present the first reinforcement learning (RL)-based solutions.

#### 2.6.1 Stackelberg Defender

The interaction between the beacon and the attacker is modeled as a Stackelberg game in which the leader moves first (attacker), and the other player (the beacon) moves subsequently, making decisions based on the leader’s query. The attacker’s strategy in our game is represented by *Q* where *Q*_*j*_ represents the index of *j*^*th*^queried SNP where 1 ≤ *j* ≤ *k* ≤ *m*. Unlike in the previous attacks, the attacker can query SNPs not carried by the victim to confuse the beacon service. ***Q*** = [*Q*_1_, *Q*_2_, … , *Q*_k_], *Q*_*j*_ ∈ {1, 2, … , *m*}, 1 ≤ *j* ≤ *k* ≤ *m*.

The beacon’s strategy in the game is represented by the vector ***h*** where *h*_*j*_ ∈ [0, 1] for the query *j* which indicates the honesty rate. We discretize ***h*** for this game and let the beacon choose from four available strategies. The granularity can be increased at the cost of computational time. ***h*** = [*h*_1_, *h*_2_, … , *h*_*k*_], *h*_*j*_ ∈ {0.25, 0.5, 0.75, 1}, 1 ≤ *j* ≤ *k* ≤ *m*.

We define the utility of the attacker as 1 − *p*_*victim*_(*Q*, **x, h**, *C*) which is the complement of the victim’s privacy. The utility of the beacon consists of two parts: (i) the privacy term and (ii) the honesty term. The privacy term first computes the LRT values of the individuals in the control group *C* for the *i*^*th*^ query using strategy *h*_*i*_. Then, it computes the variance in the first quartile of these values to quantify the risk of privacy violation. High variance indicates that an incoming query is posing a re-identification risk. The honesty term is simply *h*_*i*_. The final term is the weighted sum of these two terms where weights are hyperparameters to the algorithm. Please see Supplementary Note 1 for the details about the beacon’s utility calculation.

#### 2.6.2 Reinforcement Learning-based Attackers and Defenders

In this section, we discuss our Reinforcement Learning (RL) approach designed to protect individuals’ privacy in the beacon database while preserving system utility. The environment has two main players: the user (regular or attacker) and the beacon service.

##### States

The beacon agent’s state space *S*_*b*_ consists of the user’s *t*^*th*^ query *Q*_*t*_ and the beacon service’s statistical data at time *t*. We define ***s***_***b***_ as the beacon state, where ***s***_***b***_ ∈ *S*_b_. Specifically, each state includes (i) the MAF of the queried SNP *j*, (ii) the minimum and mean LRT *L*_*i*_(*Q*, ***x, h***) values among the individuals in each of beacon database and control group before responding the current query, (iii) potential LRT change by lying or not for SNP *j*, (iv) the minimum *p*_*i*_(*Q*, ***x, h***, *C*) among beacon participants after lying or not, and finally, (v) the beacon’s utility *u*(*Q*, ***h***) up to the current query *Q*_*t*_. Importantly, the agent only observes summary statistics rather than individual genomes which reduces state complexity and enhances privacy, especially in decentralized or outsourced training scenarios.

The attacker’s state space *S*_*a*_ contains information about the victim. The state ***s***_***a***_ ∈ *S*_a_ categorizes the victim’s SNPs into *g* groups: *g* − 1 groups based on their MAF values and another group for SNPs that she does not have. Let *s*_*min*_ and *s*_*max*_ be the SNPs with the lowest and the highest MAFs in the group, respectively, For each group, ***s***_***a***_ includes: (i) 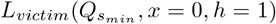, (ii) 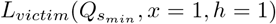, (iii) 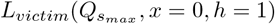, and (iv) 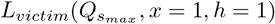. The attacker assumes the beacon is honest and calculates the LRT ranges for the rarest and most common SNP in each group as if those were the only queried SNPs. ***s***_***a***_ also contains (i) the number of SNPs, (ii) the minimum and mean LRT values obtained on the control group *C* at time *t*, and (iii) *p*_*victim*_(*Q*, ***x, h***, *C*) at time *t*.

##### Actions

The beacon agent’s continuous action space *A*_*b*_ is an honesty rate to choose for the query *j*. That is, *A*_*b*_ = {***h*** ∈ R^*k*^ | 0 ≤ *h*_*t*_ ≤ 1, ∀*t* ∈ {1, 2, … , *m*}} . The attacker’s action space *A*_*a*_ is a SNP group to choose from: *A*_*a*_ = {***a*** ∈ Z^*k*^ | 1 ≤ *a*_*t*_ ≤ *g*, ∀*t* ∈ {1, 2, … , *m*}}. From the chosen group, a specific query is then randomly selected for querying.

##### Rewards

We use both intermediate and final rewards to train our agents. The intermediate reward *r*_*t*_ is computed as a function *R* of a given state-action pair at time *t*: *r*_*t*_ = *R*(***s***_*t*_, *h*_*t*_). The function *R*_*b*_ assesses the trade-off between privacy and utility for the beacon agent for each query: *R*_*b*_ = *p*_*victim*_(*Q*, ***x***, *h, C*) + *h*_*t*_. As articulated, *p*_*victim*_ represents the privacy of the victim component, and *h*_*t*_ represents the utility of the system for *t*^*th*^ query. This ensures that each action taken by the beacon considers privacy and utility. For the attacker agent, the intermediate reward function *R*_*a*_ is defined based on the beacon’s honesty rate for the *t*^*th*^ query: *R*_*a*_ = *h*_*t*_. Here, the attacker’s goal is to maximize the honesty rate from the beacon’s answers to improve the chances of reidentifying the victim.

In the real-world scenario, the beacon does not know which individual the attacker is targeting, and the attacker is unaware of the honesty level of the responses. However, during training, information about the victim’s identity is provided to the beacon agent, and the honesty rate is provided to the attacker agent for the models to converge faster. This information is removed during inference for the system to be realistic. Additionally, the agent’s objective is to maximize a cumulative reward: The sum of all rewards that will be collected after the end of the current episode, discounted by a factor 0 *< γ <* 1 based on the time 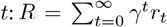. Finally, a final constant reward is given at the episode’s end based on the success of the agent: Beacon receives it if the victim is not reidentified, or the attacker earns it for reidentifying the victim.

##### Policies

The outputs of our policies are designed as computable functions that depend on a set of trainable parameters ***θ***. The beacon uses the Twin Delayed Deep Deterministic Policy Gradient (TD3) [18] algorithm which is suitable for continuous action spaces like the beacon’s. TD3 consists of two components: A policy network (actor) *µ*_*θ*_(***s***_*b*_) which selects actions *y*_*t*_ ∈ *A*_*b*_ and two networks (critics): 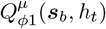 and 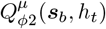, which estimate the expected cumulative reward for taking action *h*_*t*_. The critic evaluates the Q-value (*Q*^*μ*^(***s***_*b*_, *h*_*t*_)) based on the Bellman equation:

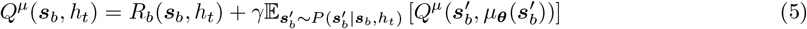

where 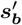 is the next state sampled from the environment’s transition model 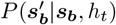. The actor is optimized to deterministically select actions that maximize the value predicted by the critic, following the objective function 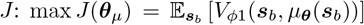. The policy *µ*_*θ*_is then updated to maximize the expected return as estimated by the critics. This update is performed by adjusting the policy parameters ***θ***_*μ*_ using gradient ascent, 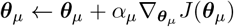 , for the policy learning rate *α*_*μ*_.

The attacker operates in a stochastic and discrete action space which requires a different approach for policy optimization: Proximal Policy Optimization (PPO) [19]. PPO also uses actor and critic networks but in contrast to TD3, (i) it uses a single critic to estimate the value function and (ii) the actor chooses actions probabilistically. We define the attacker’s policy *π*_*θ*_ given parameters *θ* and the objective function of PPO to be maximized as follows:

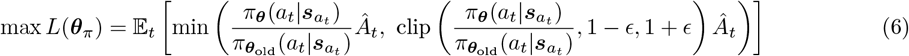

where ***θ***_*old*_ is the vector of policy parameters before the policy update, *ϵ* is the clipping term to control the learning rate, and *Â*_*t*_ is an estimator of the advantage function at timestep *t* which quantifies the benefit of choosing *a*_*t*_ over others in a given state.

We train the agents using two approaches: (1) training against predefined static strategies and (2) training against adaptive agents who can dynamically change strategies. Using approach 1, we train **the Optimal Beacon Defender (OBD)**. This agent was trained against both the optimal attacker and a set of regular users. Thus, it learns to detect the optimal attack. However, an optimal attacker may adapt its strategy by querying SNPs outside the MAF order or targeting SNPs that the victim does not carry. OBD cannot defend against this. To model this attacker, we train the **Strategic Beacon Attacker (SBA)** using approach 1, against predefined defense strategies from literature (See Section 3.3). To ensure variability and prevent overfitting, in each episode, and randomly switched the opponent’s strategy during training.

Using the approach 2, we train the **Generic Beacon Defender (GBD)** and the **Generic Beacon Attacker (GBA)** against each other in a two-player setup. The OBD and SBA are used as the starting point to initialize training. To address inherent challenges in multi-agent reinforcement learning (MARL) within adversarial environments, we implemented a structured training framework with a centralized environment and a shared control group accessible to both agents only during training, e.g., publicly available HapMap samples. Please see Section 4 for a discussion on these design choices.

## 3 Results

### 3.1 Datasets

We evaluate of our defense techniques using the 164 individuals in the CEU population of the HapMap dataset [20]. We randomly select (i) 40 individuals as the beacon participants; (ii) 50 individuals for the control group of the beacon; and (iii) 50 individuals for the control group of the attacker. For model building, we assume an attack on 10 randomly selected beacon participants. In each episode, the RL-based defense methods select one individual from this group for training, with control groups shared between the attacker and the beacon only during training. The remaining 30 beacon participants are reserved for system testing.

### 3.2 Experimental Setup

We trained OBD and SBA over 100,000 episodes, followed by a 25,000 episode fine-tuning phase for the GBD and GBA. We allowed a maximum of 100 queries per episode and performed training cycles every 10 episodes. For the beacon agent, using the TD3 algorithm, training began after 100 episodes of exploration. The agent was trained with a learning rate of 10^−4^ for both the actor and critic networks, a discount factor *γ* of 0.99, and a batch size of 256. The replay buffer size was set to 10^4^, with policy noise at 20 percent of the maximum action value and noise clipping set at 50 percent to prevent excessive exploration. The TD3 agent was trained for 50 epochs per update cycle. The attacker agent, on the other hand, was trained for 300 epochs per update cycle using a cyclic learning rate scheduler, with a base learning rate of 10^−5^ and a maximum learning rate of 10^−3^. We set the discount factor to 0.99 and applied an epsilon clipping threshold *ϵ* of 0.2 to limit policy updates. We defined the *g*=6 MAF range groups as follows: [0-0.03], (0.03,0.1], (0.1,0.2], (0.2,0.3], (0.3,0.4], and (0.4,0.5]. Training took 29 hours for single-agent sessions. Mutli-agent fine-tuning took 34 hours. The only set of hyper-parameters in the Stackelberg game is the weights for the privacy and utility terms in the defender’s reward, which we set to 0.85 and 0.15, respectively.

We evaluated all reinforcement learning models on a SuperMicro SuperServer 4029GP-TRT with 2 Intel Xeon Gold 6140 Processors (2.3GHz, 24.75M cache) and 256GB RAM. The RL agents were trained using a single NVIDIA GeForce RTX 2080 Ti GPU. The game theory approach was simulated on a SuperMicro SuperServer with 40 cores in parallel (Intel Xeon CPU E5-2650 v3 2.30 GHz).

### 3.3 Compared Methods

We compare our methods with the following baselines and state-of-the-art methods the literature. **Honest Beacon**: The defender responds truthfully to all queries, setting the lower bound for privacy and the upper bound for utility. **Baseline Method**: The defender flips *k* percent of SNVs with the lowest MAF, establishing a lower bound for effectiveness. **Random Flips**: Proposed by [4], this method randomly flips *ϵ* percent of unique SNVs, enhancing privacy by adding randomness to responses. **Query Budget**: Also from [4], this method assigns a privacy budget to each individual, limiting their exposure by reducing their contribution to responses after multiple queries. This budget decreases with each query involving the individual, especially for rare alleles, which pose a higher re-identification risk. Once the budget is exhausted, the individual is removed from future queries to preserve their privacy. **Strategic Flipping**: The Strategic Flipping [7] approach flips *k* percent of SNVs in decreasing order of their differential discriminative power, targeting the most informative variants first. The game-theoretic approach of [12] is not directly comparable to our work as it is using a non-LRT-based formulation of risk and it aims at modifying the summary statistics within the system (e.g., allele frequencies) instead of directly altering the beacon responses like we do. The code and the model were also not available.

### 3.4 Stackelberg Defender is effective yet does not scale

The Stackelberg defender’s search space expands exponentially with the number of queryable SNPs, posed queries, and potential beacon strategies, as it must evaluate all combinations to determine the optimal strategy for both parties. Refer to Supplementary Figures 1 and 2 which demonstrate that the system is infeasible for more than a few queries (*<* 6), available SNPs (*<* 8) or strategies (*<* 5).

Figure 1A shows the utility distributions for the optimal attacker and the honest beacon. Note that we use the utility definitions in Section 2.5. The distributions are obtained using the attacks against 10 victims in the beacon with 5 rarest SNPs of the victims available to the attacker. Results show that the the attacker attains a higher utility compared to the beacon as it receives honest responses for all queries posed while the beacon’s utility is much lower.

**Fig. 1:**
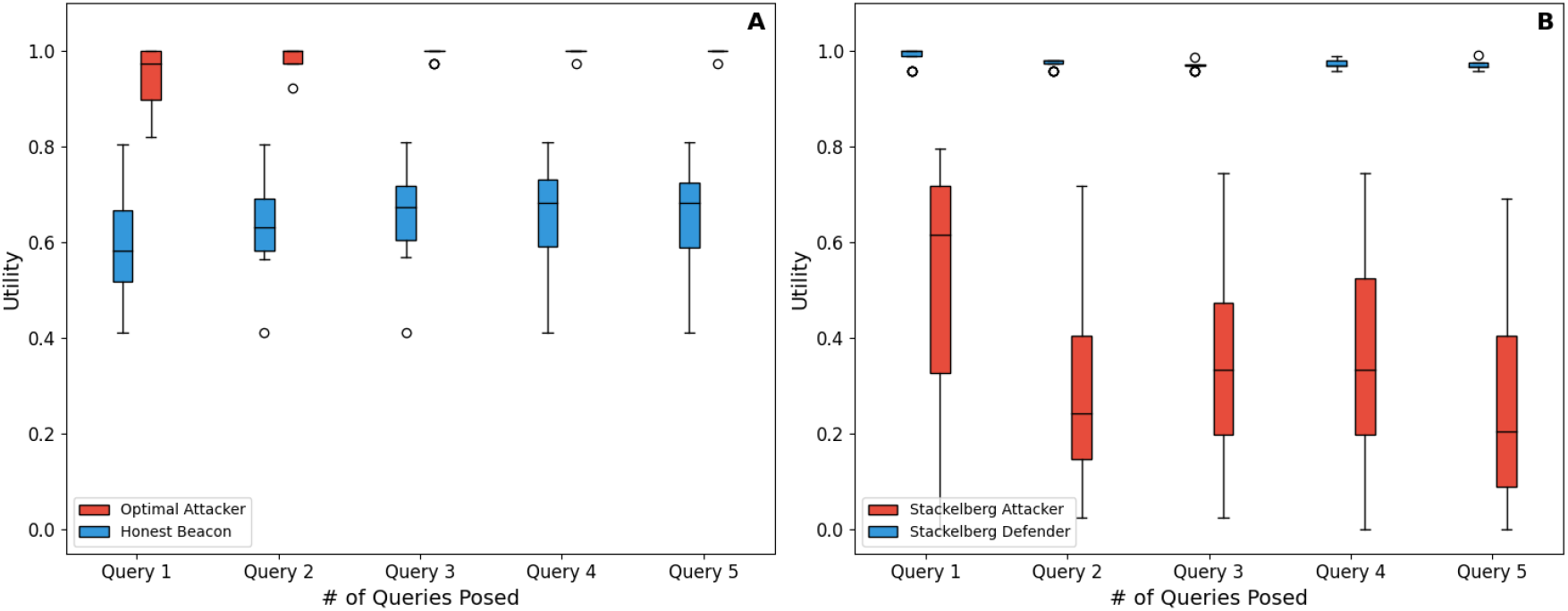
**(A)** The utilities for the *honest beacon* (blue) and the *optimal attacker* (red) across five different queries (cumulative). Each box plot represents the range of utilities achieved under each party, where the red boxes show the attacker’s utility and the blue boxes show the beacon’s utility. Outliers are indicated by circles. **(B)** The corresponding utilities in a Stackelberg game between the *Stackelberg attacker* (red) and the *Stackelberg defender* (blue) similar to **A**.

Figure 1B shows the utility distributions for the Stackelberg attacker and the Stackelberg defender playing against each other. Here, the attacker can move away from acting as the optimal attacker and can reorder the queries to mimic a regular user against the defender who can adjust her honesty level. Results show that the defender achieves much higher utility compared to the honest counterpart despite giving noisy responses. The attacker, on the other hand, cannot breach the defense and obtains a much lower utility compared to the optimal attacker despite being able to adjust its strategy. See Supplementary Figure 3 for the results when the attacker can also pose queries that the victim does not have to further confuse the defender.

We observe that modeling the beacon-attacker interaction as a Stackelberg game is effective in designing a defense mechanism, yet it does not scale beyond 5 queries and SNPs. RL-based techniques can be effective and scalable, efficiently navigating the strategy space for both parties.

### 3.5 Optimal-Beacon-Defender Agent can defend against the optimal attacker

Optimal Beacon Defender (OBD) is an agent trained to defend against the static strategy of the optimal attack alongside regular users who pose random queries. Figure 2A compares the performances of the OBD with methods from the literature as outlined in Section 3.3 up to a thousand queries. It shows that OBD outperforms all compared methods and maintains the beacon utility. The bars at the bottom of the figure shows the number of reidentified individuals out of 30 potential victims in the beacon. We observe that while maintaining system utility as defined in Section 2.5, the defense strategy does not let any individidual to be reidentified even after 1000 queries. In contrast, the next best method strategic flipping fails to protect 16 out of 30 individuals after 100 queries.

**Fig. 2:**
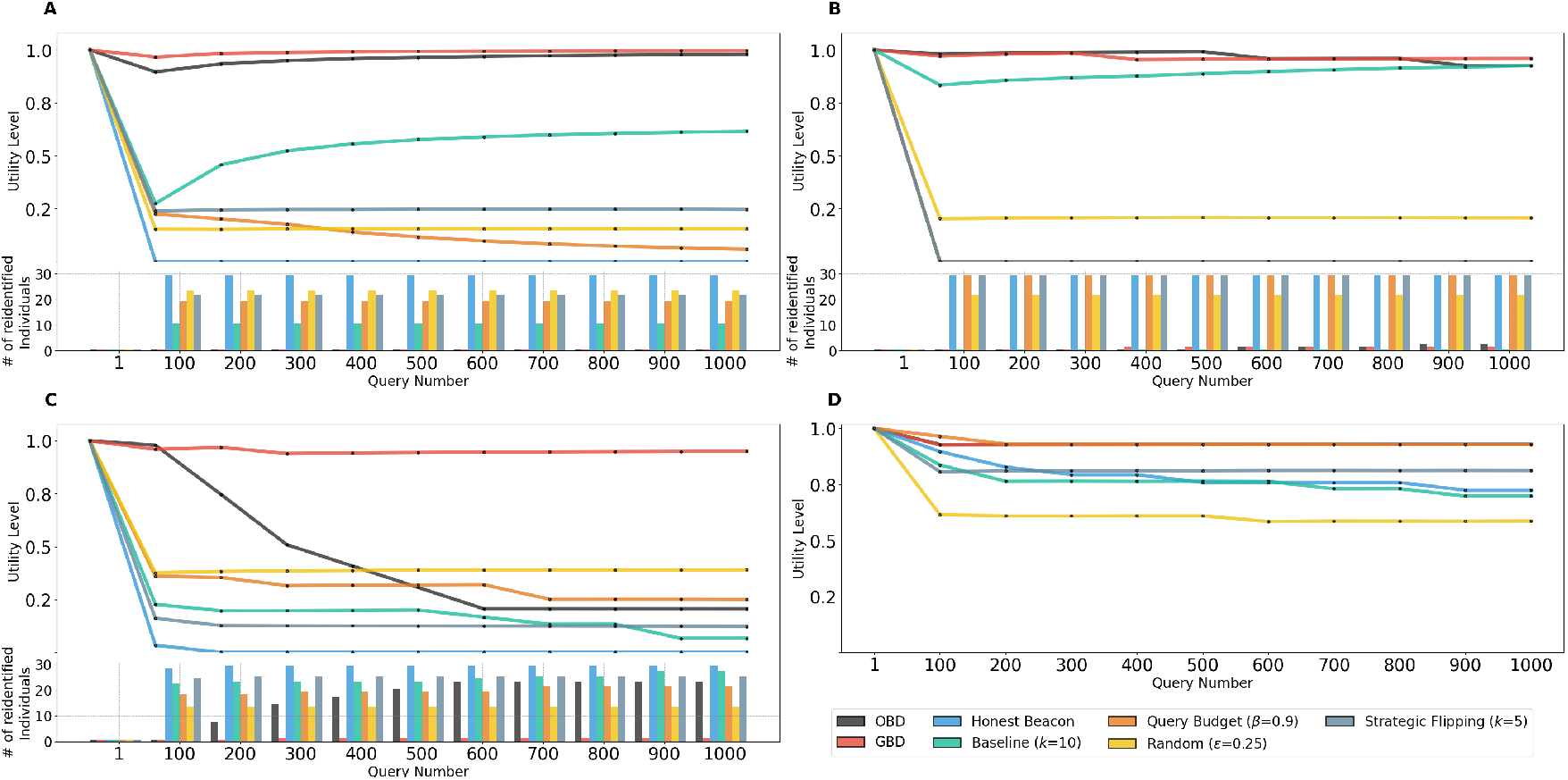
Performance comparison of Optimal-Beacon-Defender (OBD) and Generic-Beacon-Defender (GBD) against various attack strategies and query volumes. Lines indicate the utility level (y-axis) attained by the beacon (the higher the better) given a number of queries (x-axis). The bars at the bottom indicate the number of reidentified individuals at a given query volume for each defense technique (the lower the better). **A**. Performance of OBD compared to existing literature methods against the static optimal attack over 1,000 queries. OBD outperforms other methods by maintaining beacon utility and preventing reidentification for all 30 potential victims. In contrast, the next best method, strategic flipping, fails to protect 16 individuals after only 100 queries. **B**. Comparison of GBD and OBD against the SBA attacker. Over 1,000 queries, GBD maintains higher utility than OBD and prevents any reidentifications, while OBD fails to protect 2 individuals. **C**. Defense performance against the generalized GBA attacker, who can submit queries in arbitrary order and include irrelevant queries. GBD defends effectively, limiting reidentification to only 1 individual after 1,000 queries, whereas OBD fails to protect 7 individuals after 200 queries. **D**. Impact on utility for regular users submitting random queries. Both OBD and GBD fully maintain system utility, offering a significant advantage over methods that upfront add noise.

### 3.6 Generic-Beacon-Defender Agent can successfully defend against any generic attacker

Here, we test the performance of our *Generic-Beacon-Defender*, GBD, which can defend against more sophisticated attacks. First, Figure 2A shows that GBD can defend against the optimal attack and can even attain a higher utility than OBD while not letting any of the victims to be reidentified with 1000 queries. Figure 2B compares the performances of GBD and OBD against the SBA which is a more sophisticated attacker trained against other defenses from the literature. It shows that the utilities of OBD and GBD converge over time and while OBD fails to protect 2 individuals GBD does not let anyone to be reidentified over 1000 queries. Other methods attain a much lower utility and many individuals are unidentified. Figure 2C compares the performances of the defense methods against the GBA which can pose queries in any order and and can pose irrelevant queries. We observe that even OBD cannot defend against GBA and with even 200 queries 7 out of 30 people are reidentified where as GBA can defend against GBA and with 1000 queries only 1 person was reidentified. the results show that GBA can defend against any generic attacker and can dynamically adapt. Finally, we show in Figure 2D that for regular users who submit randomly selected queries, the system fully maintains the utility. This is a big advantage over methods that upfront add noise to all queries regardless of the user posing queries such as strategic flipping [7], differential privacy [9] or existing game-theoretic approaches [12]. Corresponding privacy levels across different attackers are illustrated in Supplementary Figure 5, which mirrors the subplots in Figure 2 (A, B, C, and D). This figure demonstrates that OBD and GBD achieve higher privacy level than other defense strategies, even across diverse and adaptive attack scenarios.

We also evaluate the defenses against regular users whose query selection deviates from random. We define a parameter *l* to adjust the likelihood of a user posing risky queries l ∈ [0, 1]. We define *P* (*j*) which is the probability of SNP *j* to be selected by a user given *l* as follows: such that 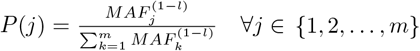. If *l* = 1 then all SNPs are uniformly selected which poses the higest risk as this increases the chances of rare SNPs to be selected. As *l* decreases rare SNPs have lower probability of being selected. We show in Supplementary Figure 4 that the beacon’s utility is very high across all considered risk groups (*l* = 0.2, 0.6 and 1) when using OBD and GBD.

## 4 Discussion & Conclusion

Multi-agent reinforcement learning faces challenges such as non-stationarity and and coordination between agents. Each agent’s learning affects the environment, complicating training. To address this, we implemented a centralized environment with consistent management of states, actions, and rewards for both agents, enhancing coordination and reducing the impact of environmental shifts caused by one agent’s learning on the other. To mitigate non-stationarity, we introduced a shared control group that both agents interact with, reducing environmental variability and providing a stable learning process. While the shared information was incorporated into the reward structure to accelerate convergence, it was not used during inference, ensuring that this mechanism aided only the learning phase and did not impact the generalization of the models. Additionally, we used separate replay buffers for each agent to store their own transitions, ensuring independent learning without interference. To prevent overfitting and ensure generalizability, we modified the evaluation population and increased the number of asked queries in evaluation tenfold compared to training, creating a more challenging test environment to assess agent robustness.

We assumed a strong adversary model which has access to the honesty rate of the responses, and yet was unable to breach the defense. Should the beacon does not release this information and follow the original protocol by just releasing the binary response, the defense would even be stronger, forcing the attacker to guess the honesty rate and adding more noise to its calculations.

A limitation of our methods is the lack of interpretability, a common issue with RL models, as we cannot clearly explain the agent’s actions unlike simpler defense mechanisms. However, this trade-off is justified by their superior performance compared to other methods. Another bottleneck is that our models are trained on a beacon composed of individuals from the CEU population, similar to many examples in the literature [3,4,5,8,6]. We demonstrate that performance with 10 training samples generalizes well to 30 unrelated individuals, allowing the model to adapt to additions or removals of participants. However, retraining is necessary if the population structure changes or for beacons with different ancestry compositions. Training takes only a few days and can be done offline. Additionally, training can be outsourced without exposing participants’ genomic data, as it relies solely on cohort summary statistics (see Section 2). In this study, we assume that a regular user submits random queries. While this assumption simplifies the modeling process, it may impose limitations in practical scenarios. In real-world applications, query patterns are likely influenced by user intent, preferences, or specific objectives, leading to non-random behavior. However, no dataset of regular user queries currently exists, restricting our ability to comprehensively evaluate the system’s robustness and performance under realistic conditions. Developing such a dataset would be highly valuable for future studies, enabling a more accurate assessment of real-world user behavior.

## References

1. GA4GH global alliance for genomics and health. https://www.ga4gh.org/about-us/. Accessed: 2024-09-29.

2. A Page, D Baker, M Bobrow, K Boycott, J Burn, S Chanock, S Donnelly, E Dove, R Durbin, S Dyke, et al. A federated ecosystem for sharing genomic, clinical data. Science, 352(6291):1278–1280, 2016.

3. Suyash S Shringarpure and Carlos D Bustamante. Privacy risks from genomic data-sharing beacons. The American Journal of Human Genetics, 97(5):631–646, 2015.

4. Jean Louis Raisaro, Florian Tramer, Zhanglong Ji, Diyue Bu, Yongan Zhao, Knox Carey, David Lloyd, Heidi Sofia, Dixie Baker, Paul Flicek, et al. Addressing beacon re-identification attacks: quantification and mitigation of privacy risks. Journal of the American Medical Informatics Association, 24(4):799–805, 2017.

5. Nora von Thenen, Erman Ayday, and A Ercument Cicek. Re-identification of individuals in genomic data-sharing beacons via allele inference. Bioinformatics, 35(3):365–371, 2018.

6. Kerem Ayoz, Erman Ayday, and A Ercument Cicek. Genome reconstruction attacks against genomic data-sharing beacons. In Proceedings on Privacy Enhancing Technologies. Privacy Enhancing Technologies Symposium, volume 2021, page 28. NIH Public Access, 2021.

7. Zhiyu Wan, Yevgeniy Vorobeychik, Murat Kantarcioglu, and Bradley Malin. Controlling the signal: Practical privacy protection of genomic data sharing through beacon services. BMC medical genomics, 10:87–100, 2017.

8. Kerem Ayoz, Miray Aysen, Erman Ayday, and A Ercument Cicek. The effect of kinship in re-identification attacks against genomic data sharing beacons. Bioinformatics, 36(Supplement_2):i903–i910, 2020.

9. Hyunghoon Cho, Sean Simmons, Ryan Kim, and Bonnie Berger. Privacy-preserving biomedical database queries with optimal privacy-utility trade-offs. Cell systems, 10(5):408–416, 2020.

10. Rajagopal Venkatesaramani, Zhiyu Wan, Bradley A Malin, and Yevgeniy Vorobeychik. Defending against membership inference attacks on beacon services. ACM Transactions on Privacy and Security, 26(3):1–32, 2023.

11. Rajagopal Venkatesaramani, Zhiyu Wan, Bradley A Malin, and Yevgeniy Vorobeychik. Enabling tradeoffs in privacy and utility in genomic data beacons and summary statistics. Genome Research, 33(7):1113–1123, 2023.

12. Tao Zhang, Rajagopal Venkatesaramani, Rajat K De, Bradley A Malin, and Yevgeniy Vorobeychik. A game-theoretic approach to privacy-utility tradeoff in sharing genomic summary statistics. arXiv preprint 2406.01811, 2024.

13. Zhiyu Wan, Yevgeniy Vorobeychik, Weiyi Xia, Yongtai Liu, Myrna Wooders, Jia Guo, Zhijun Yin, Ellen Wright Clayton, Murat Kantarcioglu, and Bradley A Malin. Using game theory to thwart multistage privacy intrusions when sharing data. Science Advances, 7(50):eabe9986, 2021.

14. Liyuan Zheng, Tanner Fiez, Zane Alumbaugh, Benjamin Chasnov, and Lillian J Ratliff. Stackelberg actor-critic: Game-theoretic reinforcement learning algorithms. In Proceedings of the AAAI conference on artificial intelligence, volume 36, pages 9217–9224, 2022.

15. Chi Cheng, Zhangqing Zhu, Bo Xin, and Chunlin Chen. A multi-agent reinforcement learning algorithm based on stackelberg game. In 2017 6th Data Driven Control and Learning Systems (DDCLS), pages 727–732. IEEE, 2017.

16. Dian Shi, Liang Li, Tomoaki Ohtsuki, Miao Pan, Zhu Han, and H Vincent Poor. Make smart decisions faster: Deciding d2d resource allocation via stackelberg game guided multi-agent deep reinforcement learning. IEEE Transactions on Mobile Computing, 21(12):4426–4438, 2021.

17. Inken Hagestedt, Yang Zhang, Mathias Humbert, Pascal Berrang, Tang Haixu, Wang XiaoFeng, and Michael Backes. Mbeacon: Privacy-preserving beacons for dna methylation data. 2019.

18. Scott Fujimoto, Herke Hoof, and David Meger. Addressing function approximation error in actor-critic methods. In International conference on machine learning, pages 1587–1596. PMLR, 2018.

19. John Schulman, Filip Wolski, Prafulla Dhariwal, Alec Radford, and Oleg Klimov. Proximal policy optimization algorithms. arXiv preprint 1707.06347, 2017.

20. Richard A Gibbs, John W Belmont, Paul Hardenbol, Thomas D Willis, Fuli L Yu, HM Yang, Lan-Yang Ch’ang, Wei Huang, Bin Liu, Yan Shen, et al. The international hapmap project. 2003.

