## Supplementary Material for "A Reinforcement Learning-based Approach for Dynamic Privacy Protection in Genomic Data Sharing Beacons"

#### 1 Supplementary Figures

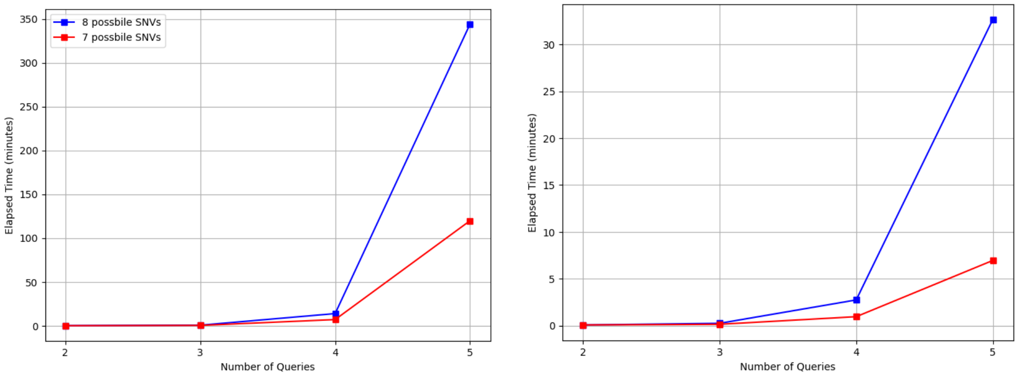


(a) (b)

Supplementary Figure 1: **a)** Execution time of the Stackelberg Game in minutes versus the number of queries for 7 and 8 possible SNPs, with the honesty rates being the same in both experiments. **b)** Execution time in minutes versus the number of queries for 3 and 4 available honesty rates, with the number of possible SNPs fixed at 6 in both experiments.

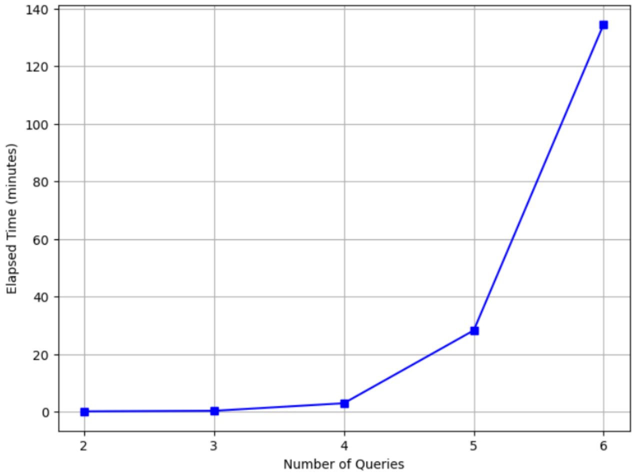


(a)

Supplementary Figure 2: Execution time of the Stackelberg Game in minutes vs number of queries for 6 possible SNPs.

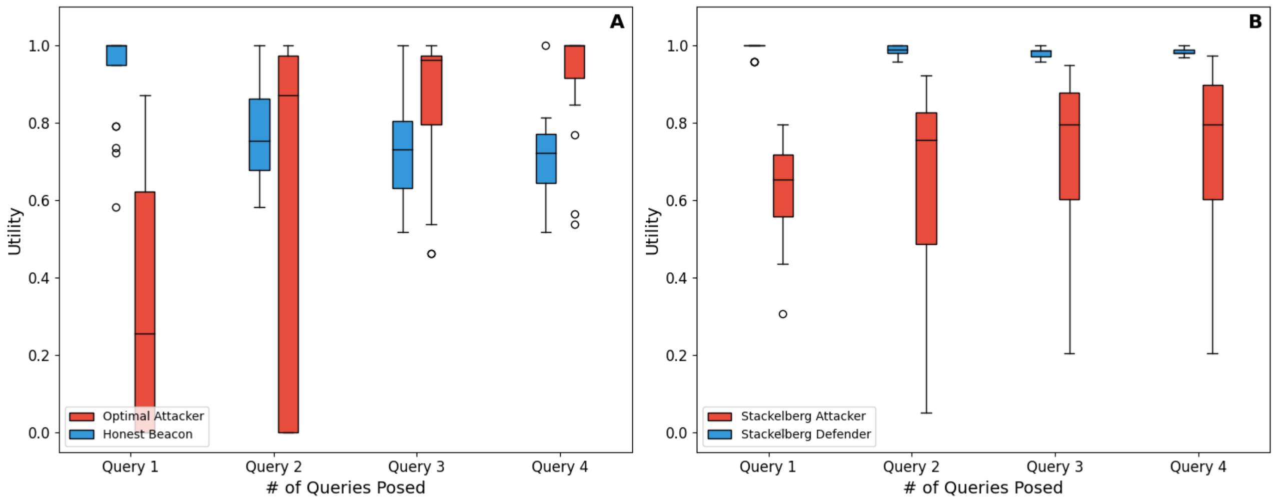


Supplementary Figure 3: **(A)** The utilities for the *honest beacon* (blue) and the *optimal attacker* (red) across five different queries (cumulative) when the attacker can query SNPs the victim does not carry to confuse the beacon. Each box plot represents the range of utilities achieved under each party, where the red boxes show the attacker's utility and the blue boxes show the beacon's utility. Outliers are indicated by circles. **(B)** The corresponding utilities in a Stackelberg game between the *Stackelberg attacker* (red) and the *Stackelberg defender* (blue) similar to **A**.

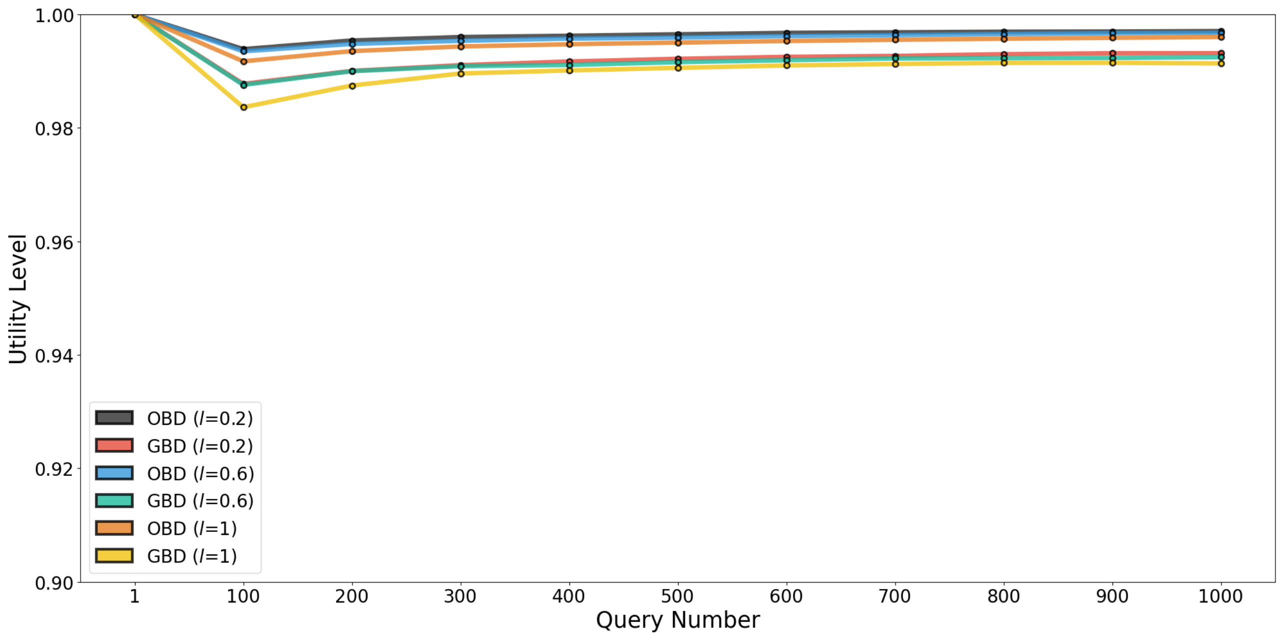


Supplementary Figure 4: Utility of the system evaluated at different risk levels ($l$ = 0.2, 0.6, and 1) for OBD and GBD models. As the risk level $l$ increases, the utility decreases for both models. The risk level parameter, ranging from 0 (low risk) to 1 (high risk), adjusts query sampling based on the Minor Allele Frequency (MAF) values. A lower risk level skews sampling towards higher MAFs (lower risk queries), whereas a higher risk level results in a more uniform distribution across all MAF values.

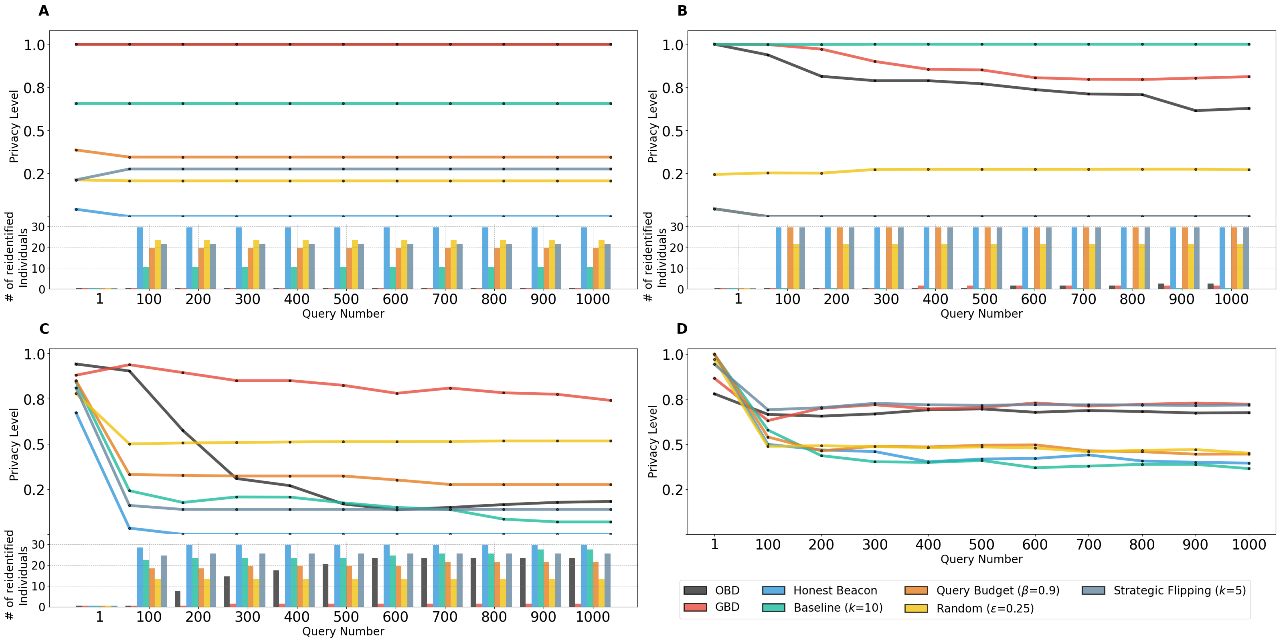


Supplementary Figure 5: Comparative performance of the Optimal-Beacon-Defender (OBD) and Generic-Beacon-Defender (GBD) across various attack types and query intensities. Privacy level (y-axis) is shown in response to increasing query volume (x-axis), with higher values indicating stronger privacy protection by the beacon. Bars below reflect the count of reidentified individuals per query volume for each defense method (lower values are preferable). **A**. Against a static optimal attack over 1,000 queries, OBD surpasses other methods by maintaining beacon privacy and preventing reidentification across all 30 potential victims. **B**. Performance of GBD vs. OBD against the SBA attacker: GBD maintains a higher privacy level and zero reidentifications over 1,000 queries, while OBD fails to safeguard 2 individuals. **C**. Defense efficacy against the GBA attacker, who poses arbitrary and irrelevant queries. GBD demonstrates resilience, allowing only 1 reidentification out of 1,000 queries, whereas OBD fails to protect 7 individuals after just 200 queries. **D**. Privacy impact for regular users submitting random queries; both OBD and GBD fully preserve system privacy, offering substantial advantage over techniques that add noise indiscriminately.

### 2 Supplementary Notes

#### 2.1 Supplementary Note 1

The algorithm to compute the beacon's utility in the Stackelberg game is as follows.

**Algorithm 1** Sharer's Utility

**function** SHARER_UTILITY($y$)<br>
    **Constants:**<br>
        *privacy_weight*, *utility_weight*<br>
    $lrt_v alues$ ← get_lrt_values()<br>
    $q_1$ ← get_first_quartile(*lrt_values*)<br>
    *target_LRTs* ← *lrt_values*[*lrt_values* $< q_1$]<br>
    **if** length(*target_LRTs*) $\neq 0$ **then**<br>
        *target_LRTs* ← append(*target_LRTs*, mean(*lrt_values*))<br>
        *target_LRTs* ← $\frac{target\_LRTs - \min(target\_LRTs)}{\max(target\_LRTs) - \min(target\_LRTs)}$<br>
    **end if**<br>
    *std_dev* ← std(*target_LRTs*)<br>
    *privacy_reward* ← $1 -$ *std_dev*<br>
    **if** length(*target_LRTs*) $= 0$ **then**<br>
        *privacy_reward* ← 1<br>
    **end if**<br>
    *privacy_reward* ← *privacy_reward* × *privacy_weight*<br>
    *utility_reward* ← [$\sum y$/length($y$)] × *utility_weight*<br>
    **return** (*privacy_reward* + *utility_reward*)/(*privacy_weight* + *utility_weight*)<br>
**end function**
